# Continuous monitoring of cost-to-go for flexible reaching control and online decisions

**DOI:** 10.1101/2022.11.16.516793

**Authors:** Antoine De Comite, Philippe Lefèvre, Frédéric Crevecoeur

## Abstract

Humans consider the parameters linked to movement goal during reaching to adjust their control strategy online. Indeed, rapid changes in target structure or disturbances interfering with their initial plan elicit rapid changes in behavior. Here, we hypothesize that these changes could result from the continuous use of a decision variable combining motor and cognitive components. We combine an optimal feedback controller with a real-time monitoring of the expected cost-to-go, which considers target- and movement-related costs, in a common theoretical framework. This model reproduces human behaviors in presence of changes in the target structure occurring during movement and of online decisions to flexibly change target following external perturbations. It also predicts that the time taken to decide to select a novel goal after a perturbation depends on the amplitude of the disturbance and on the rewards of the different options, which is a direct result of the continuous monitoring of the cost-to-go. We show that this result was present in our previously collected dataset. Together our developments point towards a continuous monitoring of the cost-to-go during reaching to update control online and make efficient decisions about movement goal.

**Author summary:** The way humans perform reaching movements is compatible with models considering that they result from the minimization of a task-related cost function. However, these models typically assume a cost function that does not change within movement, which is incompatible with experimental findings highlighting humans’ ability to adjust reaching control online and change target flexibly. We hypothesized that this later ability relied on the cost-to-go, which integrates task- and body-related information, being evaluated continuously during movement. We show that this model can optimally select and adjust control during movement in a way that reproduces human behavior in a set of tasks involving change in cost function and change in goal target. Our model predicted that decision-time to change target must be postponed when limb displacements and alternative rewards are smaller, which was borne out in our previous experimental dataset. To conclude, our model explains dynamic updates in reach control and suggests the cost-to-go as decision variable linking decision-making and motor control.

## Introduction

A commonly accepted hypothesis assumes that, when reaching towards an object, humans use a goal-directed feedback control policy tailored to the demands of the ongoing movement. This means that the observed reaching behavior is for instance tuned to the structure of the goal target (1–3), the reward associated with the movement (4–6), or the presence and location of surrounding obstacles (2,7,8). This tuning does not only characterize unperturbed movements but also the way we respond to external disturbances. In this second scenario, studies have revealed the existence of flexible feedback control loops mediated through proprioceptive (1,7,9), visual (10,11), tactile (12–14), and vestibular (15,16) sensory inputs.

Besides this ability to cope with disturbances, recent findings reported that humans respond to a change in task demands by updating their controller during an ongoing movement. This was demonstrated in previous reports by varying the structure of the target during movement, a modification which is known to elicit different control policies when these targets are presented prior to movement onset (2). Specifically, humans can exploit the redundancy of a target when there is multiple ways to attain a goal, as when reaching to an elongated object (1,2). Even when the target changed after movement onset, participants were able to alter their control policy within 150ms (17,18) through regulation of the variance and displacement along the axis that was more or less constrained according to the suddenly changing goal. Similarly, external perturbations can evoke rapid decisions to switch to a new movement goal when there are multiple alternatives. The outcome of these decisions depended on information about the target reward, the motor cost, the state of the limb, and the perturbation magnitude, which clearly points to the use of a decision variable that considers both biomechanical factors and target-related costs (19–25).

We believe that this decision variable is associated with the *cost-to-go*, defined as the total expected cost to accumulate from any point in time until the end of the movement. It plays a central role in the derivation of the optimal feedback control framework (OFC) (26,27), which posits that the closed-loop control policy underlying reaching behavior results from the minimization of the cost-function defined by a weighted sum of motor errors and control-related costs (28). Even though the OFC framework has been extensively used to explain flexible feedback control (2,29–32), it was mainly applied to static environments: that is, cost parameters have been typically considered fixed during a given trial. In light of the recent evidences for humans’ abilities to update their control policy within movements, it is necessary to enrich traditional instances of OFC models with a mechanism able to respond to these sudden changes in task parameters. On the one hand, the cost-to-go depends on the target structure and reward through the cost-function, as well as on the state of the limb including factors linked to the control of limb trajectory and potential disturbances. On the other hand, human behavioral responses to changes in movement goal described above depend on the same parameters. It is therefore natural to assume that the changes in reach behavior towards novel targets result from adjustments in control policy which reflects the dynamical estimation of the cost-to-go during movement.

To formalize these ideas, here we demonstrate that combining the OFC framework with a continuous monitoring of the task parameters, such as the target properties and the cost-to-go, captures participants’ responses to dynamic changes in the task. We modeled online modifications in control by implementing a receding horizon controller inspired by model predictive control (33,34). More specifically, at each time step, the model extracts the relevant task parameters (target structures, locations and rewards), compares the cost-to-go associated with each option when multiple are available and select the best action depending on the current task properties (e.g. target shape or lowest cost-to-go). Strikingly, this model predicts that the time of decision to switch target following an external disturbance must depend on the magnitude of the disturbance as well as on the relative rewards of the different targets. The reason is that the time required to displace the state of the limb to a region where the alternative goal becomes more attractive depends on the perturbation amplitude (smaller loads will take more time) and the relative target rewards (which modulates the size of the areas where each target is the most attractive), which is coherent with the diffusion models of decision-making (35). We reanalyzed our previous dataset and found clear evidence for a variation in decision time as predicted by the model. Altogether our developments support the theory that the brain monitors the target structure and the cost-to-go to operate dynamic and efficient adjustments of control. Our results also make a direct link between current theories of sensorimotor control and decision-making by suggesting that the cost-to-go is the decision variable used in the brain when we move in a dynamical context.

## Results

We extended the optimal feedback control framework by adding a continuous tracking of the task parameters (targets locations and structures, rewards), such that the control policy can be adjusted in dynamical contexts (changes in target structure, unexpected perturbations). We implemented a hierarchical controller, where a high-level controller adjusts the low-level Linear Quadratic Gaussian controller (LQG) to task demands, by continuously evaluating and comparing the current task parameters and potential alternative options (see Methods). For instance, when the goal target switches from a square to a rectangle during a movement, the high-level controller observes that the target shape has changed and adjusts the low-level control policy to the new target structure for the remaining part of the ongoing movement. Similarly, when the various alternative targets differ by their location and reward, the higher level of the controller compares their respective cost-to-go, which depends on task-related parameters such as target reward or the state of the system, and adjusts the low-level control policy such that the reach goal is the lowest cost-to-go target. In this implementation, the cost-to-go associated with each alternative are evaluated at each time step such that the feedback gains of the low-level controller can be adjusted to the option associated with the lowest cost-to-go. To validate this model, we reproduced a series of experimental studies involving change in target redundancy during movement and online decisions to change in the movement goal.

## Online changes in target structure

In the first behavioral study reproduced here, healthy participants were instructed to perform reaching movements to a target which could suddenly change from a narrow square to a wide rectangle or vice versa after movement started (17). Lateral step disturbances were used to elicit feedback responses and reveal potential updates in control. Participants were able to adjust the way they responded to the disturbance within movement according to the changes in target width on average 150ms after the target switched. We simulated this experiment and used the 150ms delay as a parameter to update the control policy based on the actual target structure.

In our simulations, the model captured the fact that participants let their hand deviate towards more eccentric locations along the redundant axis when the target switched from a narrow square to a wide rectangle (Figure 1A, dotted red line versus full red line), and the opposite behavior was reproduced when the target switched from a wide rectangle to a narrow square (Figure 1A, dotted blue versus full blue lines). Lateral hand deviations induced by the mechanical disturbance (Figure 1B) were similar to what was experimentally observed. Indeed, participants’ hand ended at more or less off-centered positions in the narrow-to-wide or wide-to-narrow conditions, respectively. The forward movement, aligned with the main reaching direction, was not influenced by the change in target structure while the end-point variances along the x-axis clearly depended on the condition (Figure 1C) as we reported experimentally. The transverse velocity (aligned with the x-axis) was also clearly modulated by the experimental condition (Figure 1D), which resulted from the differences in motor commands acting along that transverse direction.

**Figure 1.**
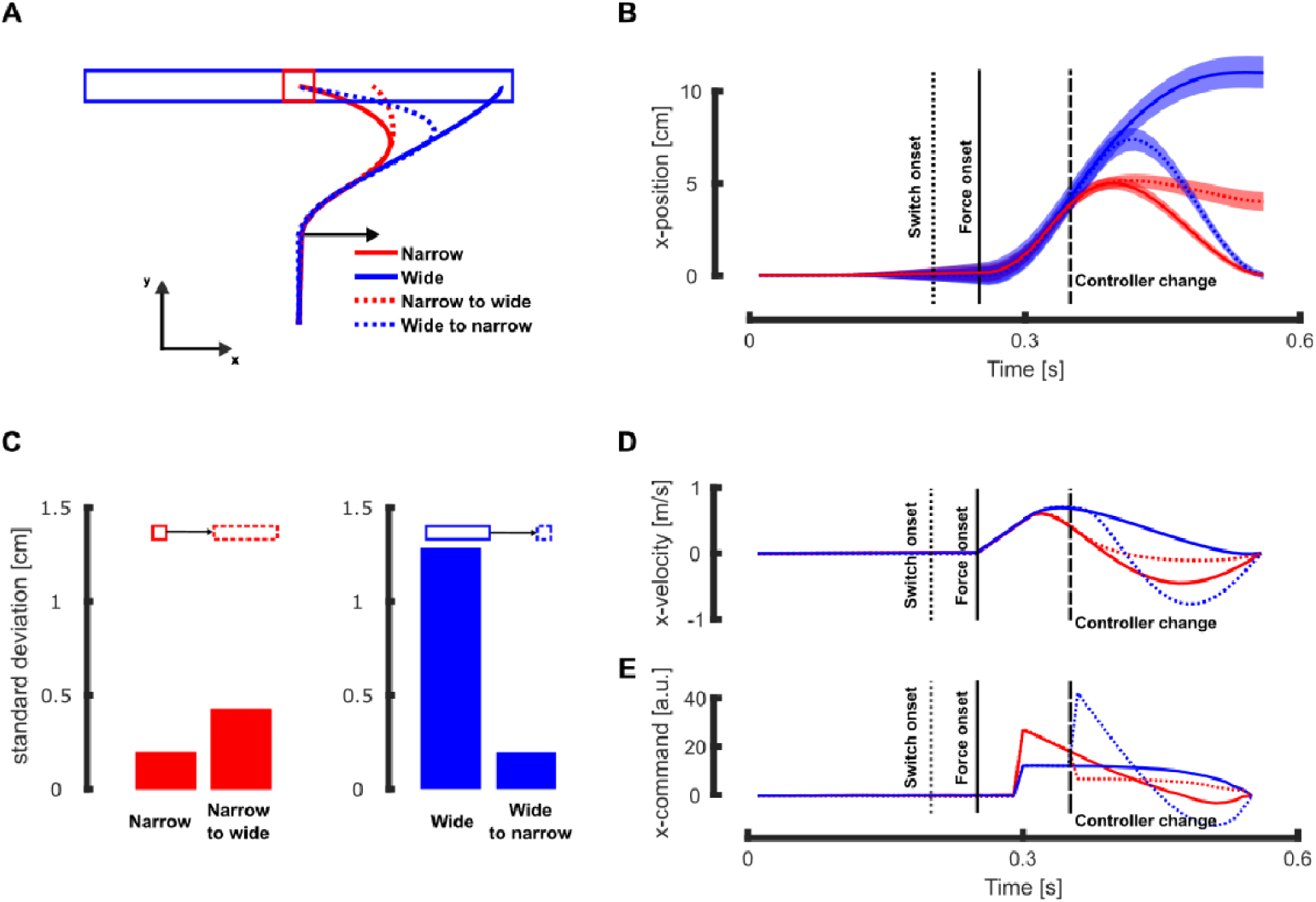
Simulation results: narrow and wide targets. **A** Mean reaching traces in presence of a rightward mechanical perturbation (represented by the black arrow) for trials initially directed towards a narrow (red) or wide (blue) target. The full and dashed lines represent the trials with and without switch in target structure, respectively. **B** Mean and standard deviation of the x-position for the different target conditions. Time is aligned with movement initiation and the vertical dotted, full, and dashed lines respectively represent the target switch onset, the force onset, and the time at which the controller was updated. **C** Simulated end-point variances along the x-axis in absence of mechanical disturbances for trials without (left bar plot) and with change in target structure (right bar plot). The left panel represents the switch from square to rectangle and the right one the switch from rectangle to square. **D** Mean traces of the transverse velocity in the different target conditions. **E** Mean traces of the x-motor command for the different target conditions.

Our model also reproduced the increase in the intensity of motor commands that can be qualitatively compared to the increase in EMG activity measured in the muscles stretched by the mechanical perturbation. Even though our implementation was not designed to fit EMG data precisely, we could compare the relative changes (increase or decrease) in motor command along the x-axis (Figure 1E) with the modulation of muscle responses following wide-to-narrow and narrow-to-wide switches. In the wide-to-narrow switch (dashed blue line), the model displayed an increase in motor command consistent with the increase in the electromyographic (EMG) activity of the stretched muscle. Similarly, in the narrow-to-wide condition (dashed red line) the model produced a decrease in motor command also compatible with our previous experimental findings (17).

Next, we confronted the model with our more recent experimental study in which we reported that online adjustments in the control policy were sensitive to dynamical factors such as a the rate of change in target width (18). For this purpose, we simulated movements towards an initially wide target (black in Figure 2A), which could continuously become narrower during movement with different rates of change. As in our experimental study, we considered three different changes in target width: a sudden switch to a narrow target (*switch* condition, magenta in Figure 2A), a fast continuous decrease in target width (*fast* condition, blue in Figure 2A), and a slow continuous decrease in target width (*slow* condition, green in Figure 2A), see Methods for the detailed implementation of these conditions within our model.

**Figure 2.**
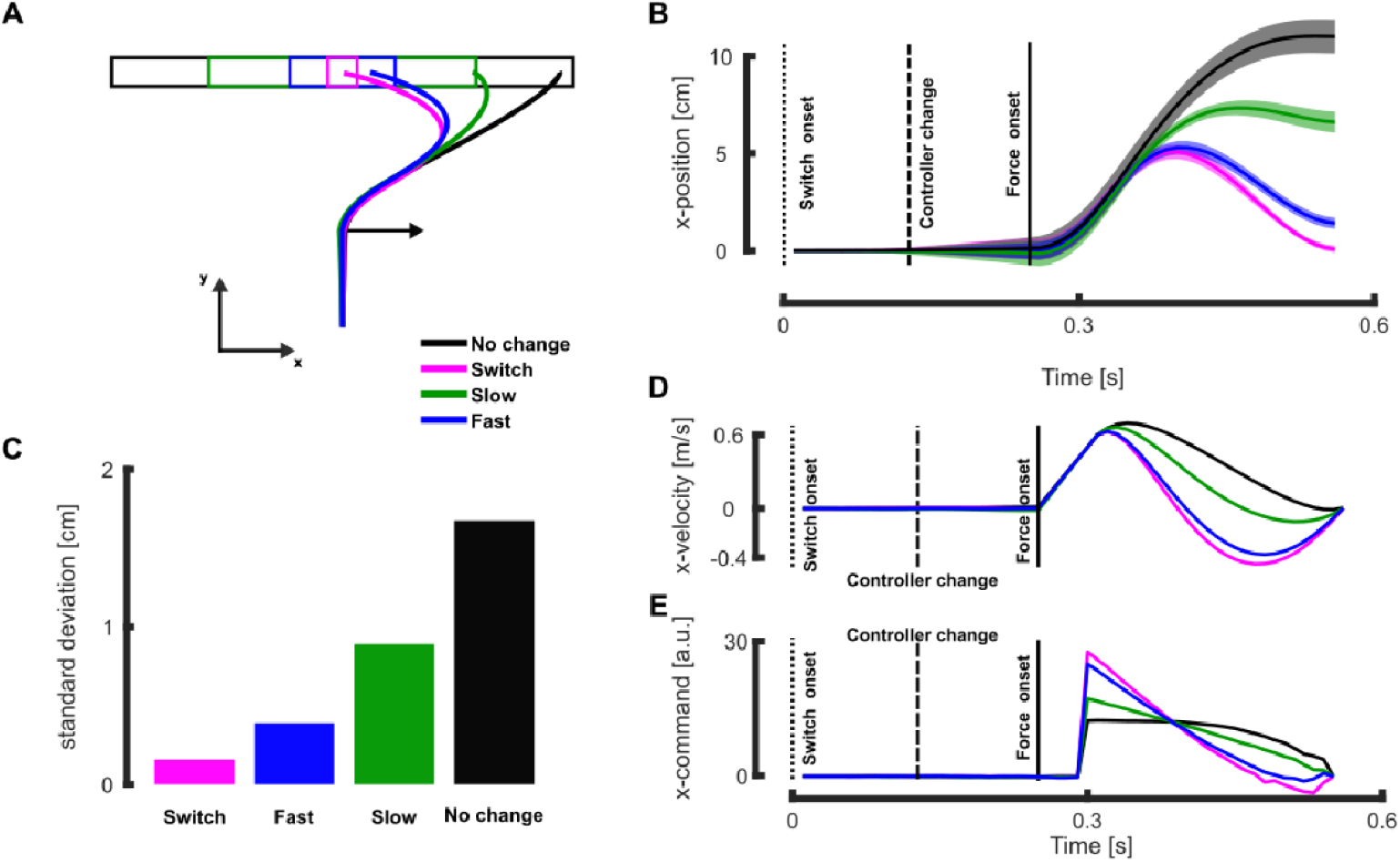
Simulation results: Continuous change in target width. **A** Mean reaching traces in presence of a rightward mechanical perturbation (represented by the black arrow) for the different target conditions in the absence (black) or presence of target change (green: slow continuous change, blue: fast continuous change, and magenta: instantaneous change). **B** Mean and standard deviation of the x-position for the different target conditions. Time is aligned with movement initiation and the vertical dotted, dashed, and full lines respectively represent the target switch onset, the time at which the controller was updated, and the force onset. **C** Simulated end-point variances along the x-axis in absence of mechanical disturbances for the different target conditions. **D** Mean traces of the transverse velocity for the different target conditions. **E** Mean traces of the x-motor command for the different target conditions.

Our model reproduced the behavioral observations for each target condition (Figure 2A). The end-point distribution along the x-axis, which indirectly quantifies how much the perturbation has been corrected, clearly depended on the target condition. The mean end-point position was more eccentric for less constraining target conditions. The model also captured how the target condition influences feedback responses to mechanical perturbation, leading to changes in lateral hand deviation and velocity along the lateral axis (resp. Figure 2B and D). Changing the target structure did not affect the movement along the main reaching direction, consistent with our experiments and the minimum intervention principle that is a known feature of human feedback control strategies (27). We also captured the patterns of activations in the stretched muscles reported in the experimental study thanks to the motor command along the x-axis. We observed that this motor command was modulated by the target condition and that it even scaled with the rate of change in target width such that the more constraining changes in target structure were associated with larger motor command (Figure 2E).

Importantly, the model captures the fact that the target structure modulates both the mean behavior and its variability. When the target width was reduced during movement, the movement variability along the transverse axis decreased as well (see Figure 1B, dotted vs full blue and Figure 2B), the mirror effect was observed when the target width increased during movement (Figure 1B, dotted vs full red). This modulation of the behavior variability is further illustrated by the end-point distribution, showing a clear dependency on the target condition for the unperturbed movements (Figure 1C and Figure 2C). This indicates that the controller fully exploited the target redundancy to control movement, similarly to what was reported experimentally.

Together, these results demonstrate that the recursive computation of feedback gains that we proposed to model tasks involving online alteration of the task demands was able to reproduce (1) online updates in the structure of the controller, (2) modulation according to dynamic factors such as the rate of change in target width, and (3) exploitation of target redundancy visible in the end-point coordinates and variances. This recursive implementation of motor control which integrates changes in the environment can be further extended to study participants’ reaching strategies in presence of multiple alternative goals as is presented below.

### Online motor decisions between alternative motor goals

Decisions to commit to a specific target are influenced by a wide range of factors, such as the reward of each target (21), the biomechanical costs (19), or even the task constraints (24). Here we formulate the hypothesis that the selection of a target (offline or during movement) directly results from an evaluation of the cost-to-go, and that online changes in movement goal occur when the estimate of the cost-to-go for an alternative target becomes more attractive than the one associated with the initial decision. Such changes can occur for instance when a perturbation pushes the hand towards an alternate goal or when the respective reward of the different targets suddenly changed during movement.

To integrate the reward of each option in the cost-to-go, we additively biased the cost of less rewarding targets with a positive constant (see Methods for more details). Biasing the cost-to-go penalized these targets and favored decisions to reach for targets with higher rewards. To validate this implementation, we simulated our previous experiment in which participants had to reach for any of three alternative targets that could be associated with different rewards (21). We show below that our implementation was able to predict the patterns of online motor decisions, in particular the statistical dependency on the frequency of switch on both the reward of the alternate target and on the perturbation magnitude (Figure 3).

**Figure 3.**
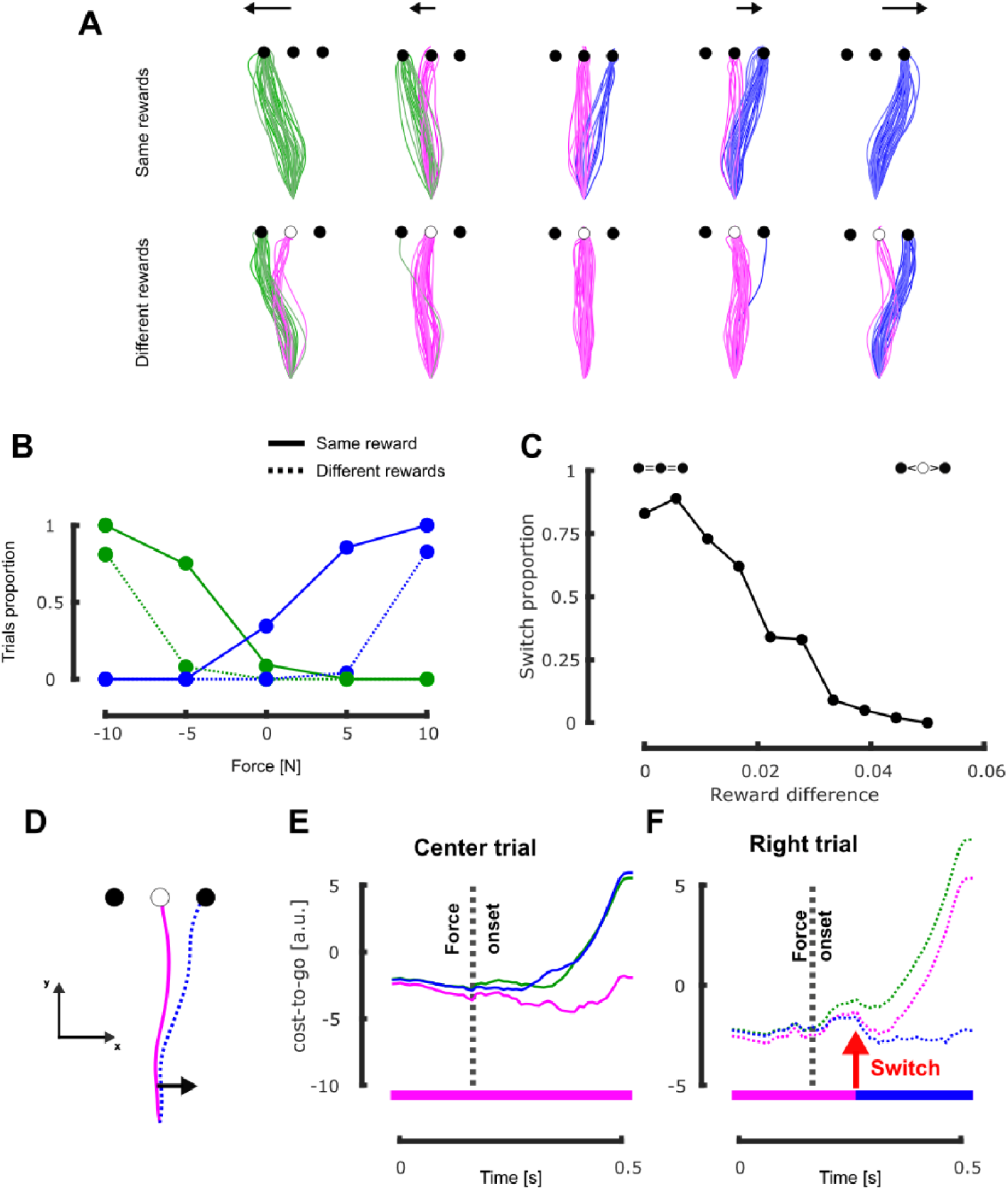
Simulation results: Online motor decisions. **A** Individual simulated hand traces in presence of multiple alternative targets (green, magenta and blue traces represent trials reaching the left, center and right target respectively). Each row corresponds to a different reward condition (top and bottom for same and different rewards, respectively) and each column represents a different force level for the mechanical disturbance (−10N, −5N, 0N, 5N, and 10N from left to right). **B** Proportion of the trials that reached the lateral targets (green and blue lines for left and right targets, respectively) as a function of the intensity of the mechanical perturbation. Full line represents the same reward condition and dotted line represents the different rewards condition. **C** Proportion of trials that reached the rightward target in presence of a rightward disturbance (5N) as a function of the difference in reward between the central and the right targets. Positive difference values favor the central target which was more rewarding than the lateral ones. **D** Hand traces for the illustrated example of the relationship between cost-to-go function and behavior, corresponding to the fourth condition of the second row of panel A (different rewards, slight rightward perturbation). The full magenta line represents a trial where participant’s hand reached the central target (corresponding to the graph of panel **E**) and the dotted blue line represents the one that reached the right target (corresponding to the graph of panel **F**). **E-F** Representation of the cost-to-go values associated with each target (green = left, magenta = center, blue = right) for trials in the different rewards condition and rightward mechanical perturbation. Panel **E** represents the cost-to-go values for the trial that ended at the central target and the panel **F** represents those for the trial that ended at the right target. The red arrow captures the time at which the right target became the goal target for the right trial and the rectangular insets at the bottom of the panels represent the target associated with the lowest cost-to-go at each time. Time axis is aligned on movement onset.

These simulations reproduced the increase in the frequency of switches to the lateral targets after the application of a mechanical disturbance when the load was larger or when the lateral target had a comparable reward (21,25). So, the model captures the behavioral observation that these two parameters played a significant role in the statistical model of target switches (Figure 3A-B). To further quantify the impact of the reward distribution on switching behaviors, we simulated a wider set of reward distributions and observed that, the larger the difference between the central and lateral targets (i.e. the more the central target was rewarding compared to the other two), the smaller the frequency of reach to the lateral targets (Figure 3C).

Importantly, the commitment to reach a target was not imposed and the controller could select to reach whichever target was more profitable according to the current estimated cost-to-go. In the experiment that we simulated, the selected target prior to the onset of mechanical perturbation was the central target as it was significantly closer to the hand than the other two (see Figure 3D-F, the magenta line lies below the other two before force onset). The mechanical perturbation altered the cost-to-go associated with each target by displacing the state of the system physically, which impacted the biomechanical factors and the cost of correcting for the perturbation, to a point where even less rewarded targets could become more profitable in comparison with the larger reward of the center target that requires a larger and costlier corrective action (see the small bump at the dashed line in Figure 3E). In the trial illustrated in Figure 3F, the controller selected a different target with lower cost-to-go value (the red arrow in Figure 3F captures the switch to the right target), which was less rewarding but also less effortful.

The implementation we proposed to simulate online motor decisions can also be used to simulate movements towards a redundant target such as a rectangular target. To show this, we extended our simulations to other experimental protocols investigating online motor decisions towards redundant targets associated with a non-uniform reward distribution along the redundant axis (20,23). In their recent study, Cos and colleagues instructed participants to perform reaching movements towards a rectangular target that had non-uniform reward distributions (20). We simulated a similar experiment and considered three different cases for the reward distribution, as in the original experiment. In the first case, the reward distribution was symmetric with respect to the center of the target (Figure 4A, Case 1). The two other cases considered asymmetric reward distributions with either a leftward (Figure 4A, Case 2) or a rightward (Figure 4A, Case 3) bias.

**Figure 4.**
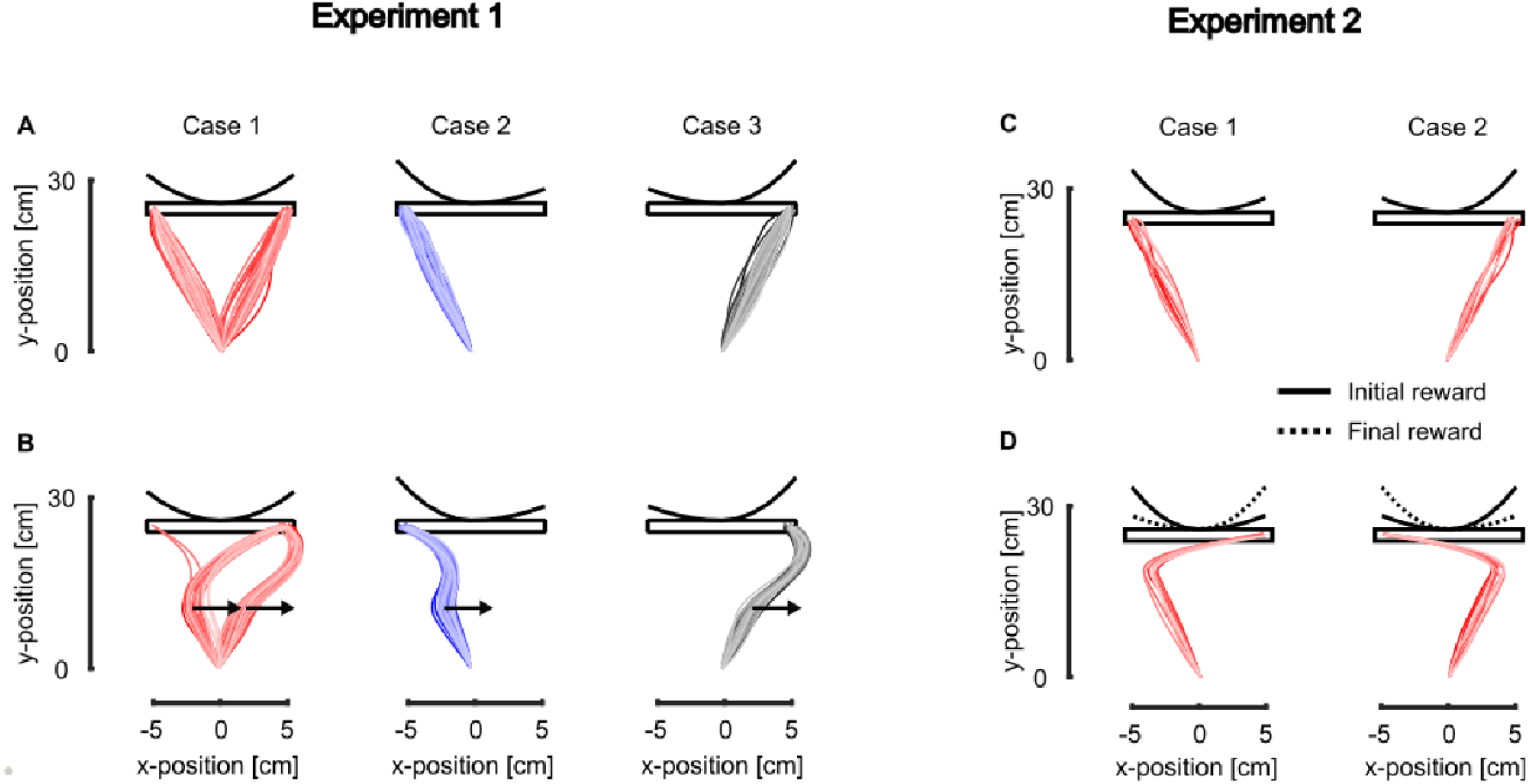
Online motor decisions for rectangular targets. **A** Individual simulated hand traces for the symmetric (Case 1) and asymmetric (left bias and right bias for cases 2 and 3, respectively) reward distributions. The reward distributions along the x-axis are represented above the targets. **B** Individual simulated hand traces for the three different reward distributions in presence of a rightward mechanical perturbation. **C** Individual simulated hand traces for the biased distributions (left and right bias represented by cases 1 and 2, respectively). **D** Individual simulated hand traces in presence of a switch in the reward distribution (from the full line to the dotted line) for both initial reward distributions.

Our model captured participants’ reaching behavior in both unperturbed and perturbed conditions. In the unperturbed condition, the model predicted that participants were biased towards locations associated with the highest reward: both extremities on the target in the symmetric condition (Figure 4A, Case 1) or the extremity associated with the largest reward in the asymmetric conditions (A, Cases 2 and 3). Interestingly, it also predicted the *changes of mind* observed in the original study by Cos and colleagues as can be observed in the symmetric reward conditions in presence of rightward mechanical perturbation (Figure 4B, Case 1), where some trials initially directed to the left end of the target ended up at the right end after the application of the perturbation.

We can combine the bias of the cost-to-go function with an online change in the reward distribution itself to model human behavior in tasks involving online change in task demands and online motor decisions between multiple targets. In a recent study, Marti-Marca and colleagues (23) instructed participants to reach for a target represented as a wide rectangle aligned with the x-axis which has a non-uniform reward distribution along this axis. In the present study, we only considered the reward distributions schematically represented above the targets in Figure 4C. As participants were reaching towards this target, the reward distribution could suddenly switch from one condition to the other (i.e. the reward bias moved from left to right or vice-versa). Our model predicted that the online changes in reward distribution impacted the reaching behavior similarly to what has been reported experimentally (23). Indeed, in Figure 4D we reported that the trials that were initially targeted towards the end of the target associated with the high initial reward (full line traces in Figure 4 D) suddenly redirected towards the other end after the reward distribution was changed (dotted line traces in Figure 4 D). This result further demonstrates the ability of our model to predict the outcome of online motor decisions in dynamic contexts.

### Experimental Evidence for Continuous Monitoring

Our strongest claim is that the nervous system monitors the cost-to-go in a continuous fashion. The alternative hypothesis is clearly that the evaluation of the cost-to-go that underlies motor decisions is not continuous. We know that decisions to switch target depend on the occurrence of an external perturbation, thus the minimal candidate model that does not feature a continuous monitoring of the cost-to-go is a discrete evaluation of this quantity, potentially triggered by the occurrence of the load disturbance. There is a testable difference between such an event-triggered switch and a continuous monitoring: in the case of a discrete switch, for a given perturbation magnitude, the decision must be taken at the same time irrespective of the different target rewards. In contrast, the continuous model features the possibility that decisions are taken at different times. The reason is that, in the condition where the lateral target has lower reward, it requires a larger hand displacement before it becomes more attractive, thus the decision to switch target is postponed until the hand has travel a larger distance, if it does. Similarly, for different perturbation magnitudes and a fixed reward, the same reasoning suggests that decisions to switch can be taken faster following larger perturbation magnitudes. Thus, contrary to a discrete switch hypothesis without ad hoc tweaking of the decision time, a continuous monitoring implies that decisions can be taken at different times as a consequence of the continuous evolution of the limb trajectory.

This effect was observed both in simulations, as expected, and in our previous dataset. In the simulations a clear delay in decision time was reproduced when lowering the reward of the lateral target gradually (Figure 5A). In addition, we reanalyzed our previously published data in search for a delaying of the decision time dependent on target reward and perturbation magnitude. We compared for each combination of perturbation direction and reward distributions the time at which trials reaching the central and lateral targets could be distinguished. We used the receiver-operator characteristic (ROC) technique to determine when these trials could be distinguished (see Methods). We found that this time onset was modulated across reward conditions and perturbation directions (Figure 5B). Strikingly, we observed, exactly as in the model simulations, that the onset of decision occurred earlier when all the targets had the same reward (leftward perturbations 147 vs 190ms Figure 5C vs D and rightward perturbations 163 vs 186ms) than when the central target was more rewarding. Similarly, we found in the same dataset that the amplitude of the perturbation influenced the decision time: smaller perturbation amplitude resulted in longer decision time (0.161s (small) vs 0.147s (large) for the leftward perturbations, Figure 5E vs F, and 0.191s (small) vs 0.163s (large) for the rightward perturbations). Even if our model was able to qualitatively predict the modulation of decision time with the conditions, the predicted time does not match the one from the data probably because the model does not implement all the mechanisms underlying the control of movement such as short-latency reflexes for instance. We indeed designed the model such that it can qualitatively, rather than quantitatively, reproduce and predict experimental results (see the standardized parameter design in Methods).

**Figure 5.**
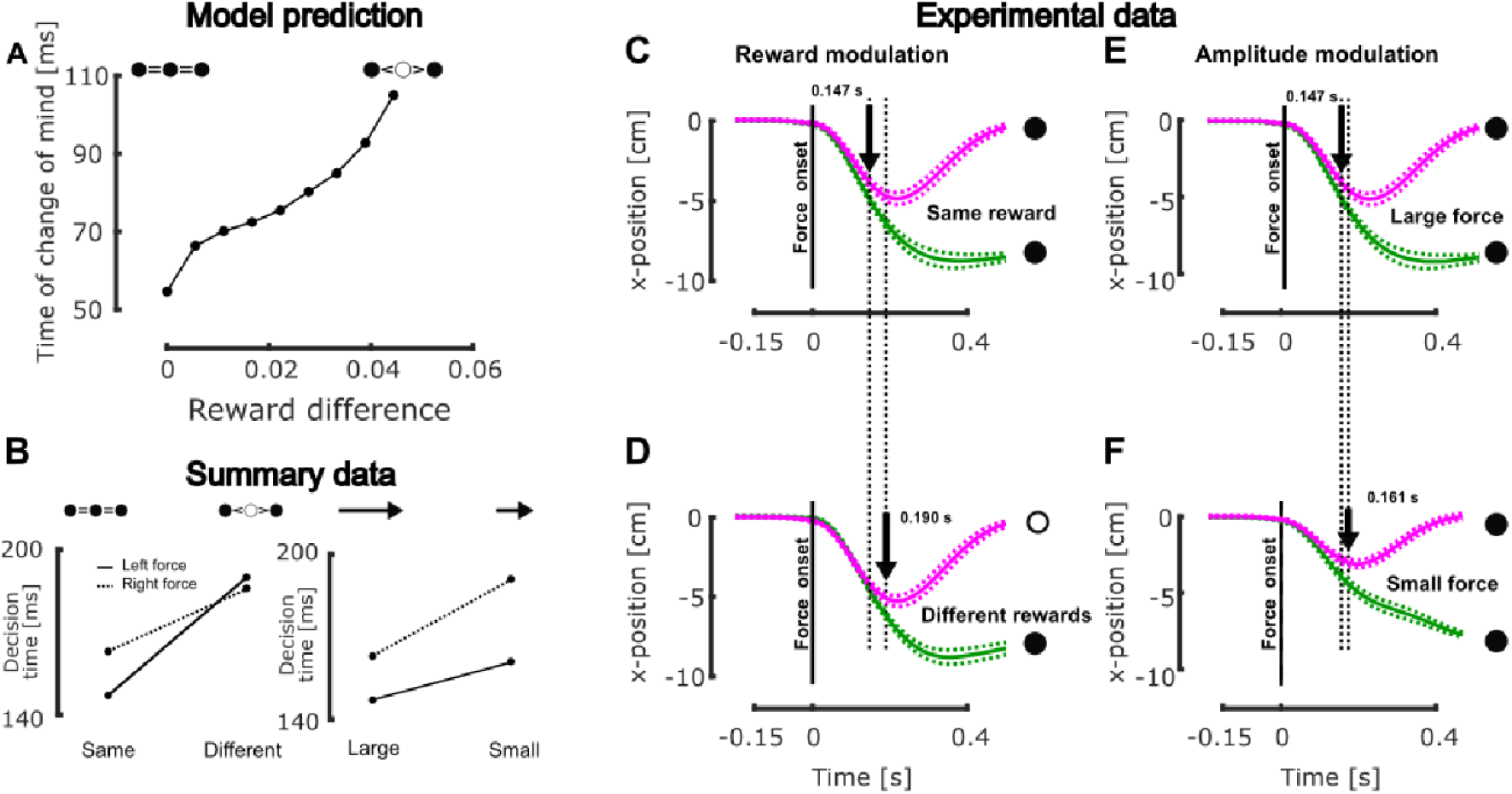
Behavioral evidence of the modulation of decision time. **A** Model prediction of the modulation of the time of change of mind as a function of the difference between the central and the lateral targets. In presence of larger reward difference (when the central target has a larger reward than the other two) the model predicted longer decision time. **B** Summary of the experimentally measured decision time for constant force and various reward distributions (left) and constant reward distribution and different force levels (right). The full and dashed lines represent the trial with a leftward and rightward perturbation, respectively. **C** Group mean and SEM hand traces along the x-axis for reaching performed in presence of three alternative targets and leftward perturbations. All the targets had the same reward, the magenta trace captures trials that reached the center target while the green trace captures those that reached the left one. The vertical dashed line represents the onset of mechanical perturbation and the black arrow the onset of change of noticeable differences between the two targets. **D** Same as **C** in the condition where the central target was more rewarding than the lateral ones. **E** Same as **C**. **F** Same as **E** in presence of small leftward perturbations instead of large ones.

In all, these results demonstrate that our hierarchical implementation of OFC which leverages the cost-to-go function is able to capture (1) reaching behavior in presence of online changes in target structure and (2) the decision to switch to an alternative reach goal in presence of change in task in a way that depends on the perturbation and reward distributions. Importantly, our model predicted that the time required to switch to a novel goal should depend on the amplitude of the perturbation and on the reward distribution since the cost-to-go was defined based on the external target reward and the time-varying state of the limb. This prediction was verified by reanalyzing our previous dataset.

## Discussion

In the present work, we tested whether human reaching behavior in dynamical environments could be captured by an optimal feedback control (OFC) model modified to continuously track the target structure and the cost-to-go of current and alternative goals, enabling online adjustments in control as well as decisions to reroute the ongoing movement to a novel goal. We implemented a recursive controller which was able to reproduce online motor decisions that depend on the state of the system, the presence of external disturbances and the respective reward of the different available reach options. It also prompted us to reanalyze previous data in search for a dependency of the decision time on target reward that was waiting to be discovered.

Clearly, the update in control that accompanies sudden changes in goal requires to change the controller online. To accommodate the experimental observations made about the dependency of these changes, we proposed an extended implementation of the OFC framework combined with a decision module based on a continuous observation of the target structure and an estimation of the cost-to-go function. Here, the idea of hierarchical control is used to refer to the organization of the algorithm, which sequentially monitors the task requirements (target structure, reward) then selects the motor command according to the best option. The concept of hierarchical control was previously used in motor control to cope with the curse of dimensionality inherent to the control of a neuromuscular model of the human body (36–38) and to bring together control and adaptation during reaching and locomotion (39,40). The present work innovates in that we exploited this hierarchical control to reproduce within-movements updates in control policy. More specifically, the lower level of our model implements a classical Linear Quadratic Gaussian (LQG) controller, often used to simulate reaching movements (2,27), and the higher level combines the notions of Model Predictive Control (MPC) and estimates of the cost-to-go to evaluate and compare the concurrent options and select the best one.

The MPC framework, which assumes that the control problem is recursively solved during movement (34,41), has already been used to explain experimental features that were not captured by other models such as the temporal evolution of feedback gains during reaching movement (42) or the modulation of the time horizon of reaching movements in presence of perturbations (33,43,44). In a recent work, this framework was combined with an impedance controller to demonstrate its ability to handle nonlinear dynamics and changing environments (i.e. force fields) during movement (45). Here, we combined this MPC framework with the continuous evaluation of the cost-to-go function by generalizing the previous implementation (25) proposed by Nashed and colleagues by integrating the reward within the cost-to-go function through a positive cost bias for the less rewarding options. Our implementation is dynamical in the sense that it computes and compares the cost-to-go of the different options at each time step in order to predict the switch in behavior at any time. This allows us to reproduce the shift in behavior and decision processes reported in multiple experimental studies involving reward (20,21,23).

In the presence of higher reward, participants will be more prone to take actions associated with higher motor costs which are for instance characterized by higher velocities and feedback responses (6,21,46).

Our main contribution is to provide support for the hypothesis that the cost-to-go is monitored continuously, which allows formulating a dynamical implementation of the distributed consensus model during movement (47). The basic premise of this model is that decisions between multiple motor actions result from the integration of high-level factors and low-level biomechanical costs, both biasing the competition with reciprocal interactions during the process leading to a motor decision (19,48). In other words, the decision to take an action depends not only on the outcome of this action (e.g. its associated reward) but also on the cost of selecting this action (e.g. its biomechanical cost). The cost-to-go function encapsulates both action outcome and motor cost. Importantly, the fact that a decision variable depends on these components was a constraint dictated by previous experimental work. Interestingly, the concomitant influence of reward and cost parameters was not only observed in tasks involving distinct reach options (Figure 3) but also in tasks that involved a redundant target (Figures 4). This does not require any adjustment of the algorithm since the cost-to-go directly follows from the cost matrices that feature or not a redundant dimension in movement goal. The model is therefore quite general, and flexible enough to handle a broad range of parameters linked to movement control.

By introducing the cost-to-go as a decision variable, we bridged an important gap between current models of motor control and decision making. We used the cost-to-go as a decision variable because of its theoretical grounding in the dynamic programming solution of the optimal control problem (49). It has been used to derive locally and globally optimal solutions to the control problem through stochastic optimal control (27,50) and reinforcement learning (51,52). The implementation presented in this paper allows to go beyond standard static formulation of OFC models with fixed parameters, and explicitly integrate the reward, changes in target structure and multiple targets in the cost-to-go, similarly to what was done with the value function (i.e. the opposite of the cost-to-go) in reinforcement learning (53).

The dynamical tracking of the cost-to-go enabled the reproduction of the behavior of healthy adults reported in multiple previous studies (17,18,20,21,23–25). The neural structure supporting these within-movement adjustments has not been experimentally identified, but we hypothesized that the basal ganglia are likely to be involved. Indeed, this structure is often associated with the evaluation of the cost of a movement (54–56) which results in the invigoration of movements often associated with the presence of reward (6,21,46,57). It is conceivable that this structure is also involved in the representation of the structure of the goal as well as in the target selection. Besides their potential implication in the evaluation of movement costs, the basal ganglia also play a central role during decision-making. Indeed, seminal models suggest that the basal ganglia compare and select the actions to be performed (58,59). The exact role of the basal ganglia in decision-making is nonetheless not yet fully understood (56,60,61). Their implication in the online adjustments process we reported in the present work are likely to be combined with other brain areas (62–65).

Further modeling works could piece together our present results with cognitive aspects of reaching control. For instance, we did not model the documented impact of reward on the modulation movement velocity and feedback gains with increasing reward (6,21,46,57,66). Previous reports attributed this modulation of movement vigor with reward to the value of time and its influence on the perception of reward (67–69). The notions of value of time and reward were integrated with optimal control in studies that confirmed this hypothesis (70–72). Recently, similar influence on reaching behavior has been reported in presence of uncertainties about the environment (73,74) which can be modeled thanks to the use of a robust controller. To account for the modulation of movement vigor related to temporal factor or to the use of a robust strategy, a fuller implementation requires an additional module which not only selects the current target or controller, but also modifies the control gains according to the vigor or robustness of the intended movement.

Together our modeling results expand the applicability of optimal feedback control framework to tasks involving dynamic changes in parameters during movement. By bridging together motor control and decision-making, the present work opens new modeling and experimental perspectives to investigate movement control in complex environments. From a normative perspective, control models must be enriched with online estimate of movement costs and updates in control. From the perspective of neural implementation, it implies that humans do not sequentially decide, plan, then act. Instead, these operations seem imbricated in a general, continuously evolving process. Finally, we believe that the similarities between the theoretical framework in which we grounded our model and that of reinforcement learning suggest direct theoretical developments to further challenge our understanding of human sensorimotor control in the context of artificial systems designed to mimic human behavior: not only reward signals must be used to train a controller but control signals must also be used to train and form an estimate of the expected reward. Exploring how this can be done in neural systems is an exciting challenge for prospective studies.

## Methods

### General model

We considered the translation of a unit point mass (m = 1kg) in the horizontal plane in presence of three forces: a viscous force proportional and opposite in sign to the velocity, a controlled force *F*, and an external force *F*_*ext*_. The controlled force was represented by a first-order low pass response of the control vector to approximate non-linear muscle dynamics. The controlled system was described by the following set of continuous differential equations that characterized movement along the x- and y-axes independently

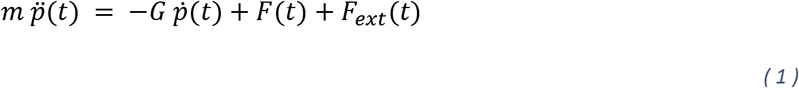

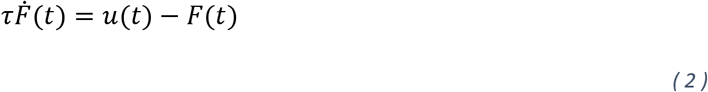

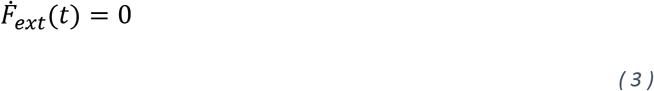

where *m* is the mass, *p*(*t*) is the two-dimensional vector of the point mass coordinate in the plane, *G* is a viscous constant, *F* and *F*_*ext*_ are the two-dimensional controlled and external forces respectively, and *u* is the two-dimensional control vector. The time constant of the linear muscle model was set to *τ* = 60ms (75). The parameters *m* and *G* were arbitrarily set to 1 to standardize the simulations. The external forces *F*_*ext*_ are assumed to be constant (Equation 3), such that any change in this variable used to simulate an external disturbance is treated by the controller as a step function.

The system dynamics was discretized using Euler method with a time step of 10ms to integrate stochasticity, which gave the following difference equation

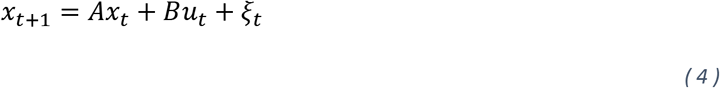

where *x*_*t*_ is the time-dependent state vector containing position, velocity, controlled and external forces, *A* and *B* are two real-valued matrices that capture system dynamics, *u*_*t*_ is the command vector at time *t*, and *ξ*_*t*_~*N*(0, Σ_*m*_) is additive motor noise defined by a multivariate normal distribution of zero mean and covariance matrix Σ_*m*_. To include the goal target, we augmented the state vector with another vector of the same dimension (dimension 16 in total, it facilitates simulations of tasks in which any subset of state variables has to be penalized). Finally, the state vector was further augmented 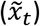 to integrate feedback delays and let the controller observe only the most delayed state (31,76). The corresponding feedback signals at each time step write

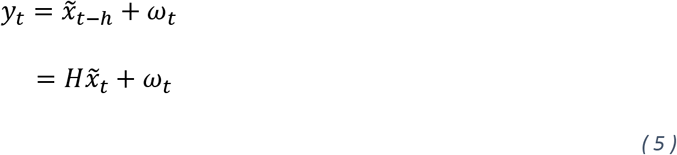

where *h* is the feedback delay, *ω*_*t*_~*N*(0, Σ_*s*_) is additive, zero-mean sensory noise with covariance matrix Σ_*s*_ standard deviation, and *H* = [0^16*x*16^, …, 0^16*x*16^, *I*^16*x*16^]^*T*^ is the observability matrix. The feedback delay was set to 50ms (h=5) to capture the sensory motor delay associated with long-latency feedback responses, assuming that this transcortical loop supports goal-directed, state-feedback control (77). Because of the noise and sensory delay that corrupt the sensory feedback, we used a dynamic Bayesian estimator weighting sensory feedbacks and priors to estimate the state vector. The maximum likelihood estimator of the posterior distribution computed through a Kalman filter is denoted 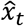 and follows

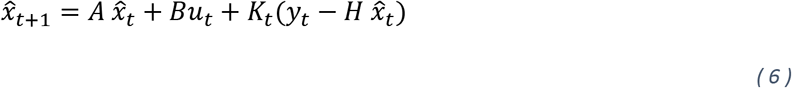

where *K*_*t*_ are the Kalman gains recursively computed (see (27) for more details). The optimal motor commands are computed by minimizing a cost-function consisting of a weighted sum of quadratic penalties on the state vector (first term) and on the motor command (second term) which writes as follows

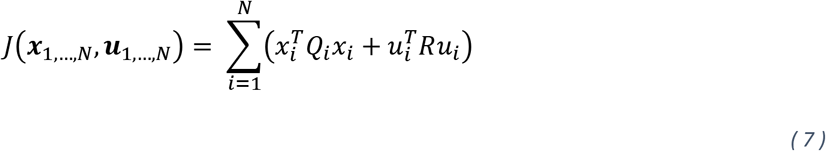

where *Q*_*i*_ ≥ 0 and *R* > 0 are real-valued matrices respectively capturing the penalties on the state vector and motor command, ***x***_1,…,*N*_ and ***u***_1,…,*N*_ capture the successive state and command vectors, and *N* is the number of time steps (i.e. time horizon). In most previous studies as well as in the present one, *N* is considered to be a finite integer which characterizes a finite horizon formulation of the control problem. Under these assumptions, the optimal motor commands 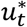 are defined by a linear combination of the different entries of the state estimate

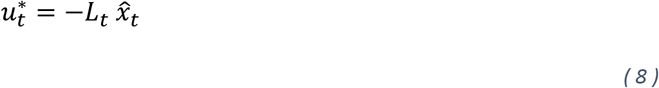

where *L*_*t*_ are the optimal feedback gains computed recursively (see (27) for more details).

### Online changes in target structure

The optimal feedback control framework allows to determine the optimal control policy for a given set of task parameters Θ = {*Q*, *R*, *N*}, capturing the structure of goal target and movement duration. The control policy obtained this way is specifically tuned to these task parameters and can be written as follows

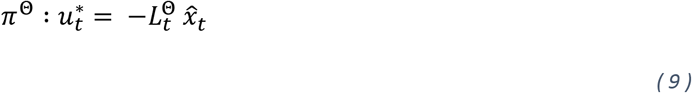

where the superscript Θ indicates that this control policy has been computed for the set of task parameters Θ. The optimal feedback gains 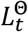 are computed offline and applied during movement execution. If we consider a second set of task parameters 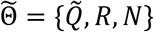 that only differs by the penalty on the state vector 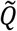, it will define an optimal control policy 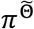 such that 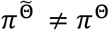. This principle has been applied in previous work to model trial-to-trial variation in task demands captured for instance by target of different shapes or different penalties to state variables (2,78). As such, this framework does not model movements in which the set of task parameters Θ, used to determine the control policy, changed during movement. However, recent behavioral experiments have demonstrated humans’ ability to optimally adjust their control policy during movements following dynamic changes in target structure (17,18).

In the case of time-varying task requirements (e.g. a change in target structure during movement), captured by a time-varying sets of task parameters Θ, we propose a recursive dynamical computation of the motor commands that considers, at any time during movement, the set of task parameters available to the controller. The basic premise of this implementation is that the set of optimal feedback control gains 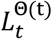 are computed at each time step from the time-varying set of task parameters Θ(t) = {*Q*(*t*), *R*, *N*} and only the first motor command is applied, similarly to what is done in the MPC framework (41). The procedure is then repeated at the next time step, for which *Q*(*t*) and *N* are adapted to respectively capture time-varying task demands and changes in movement horizon. This recursive implementation of the optimal feedback gains is schematically represented in Figure 6A, in the more general framework of OFC.

**Figure 6.**
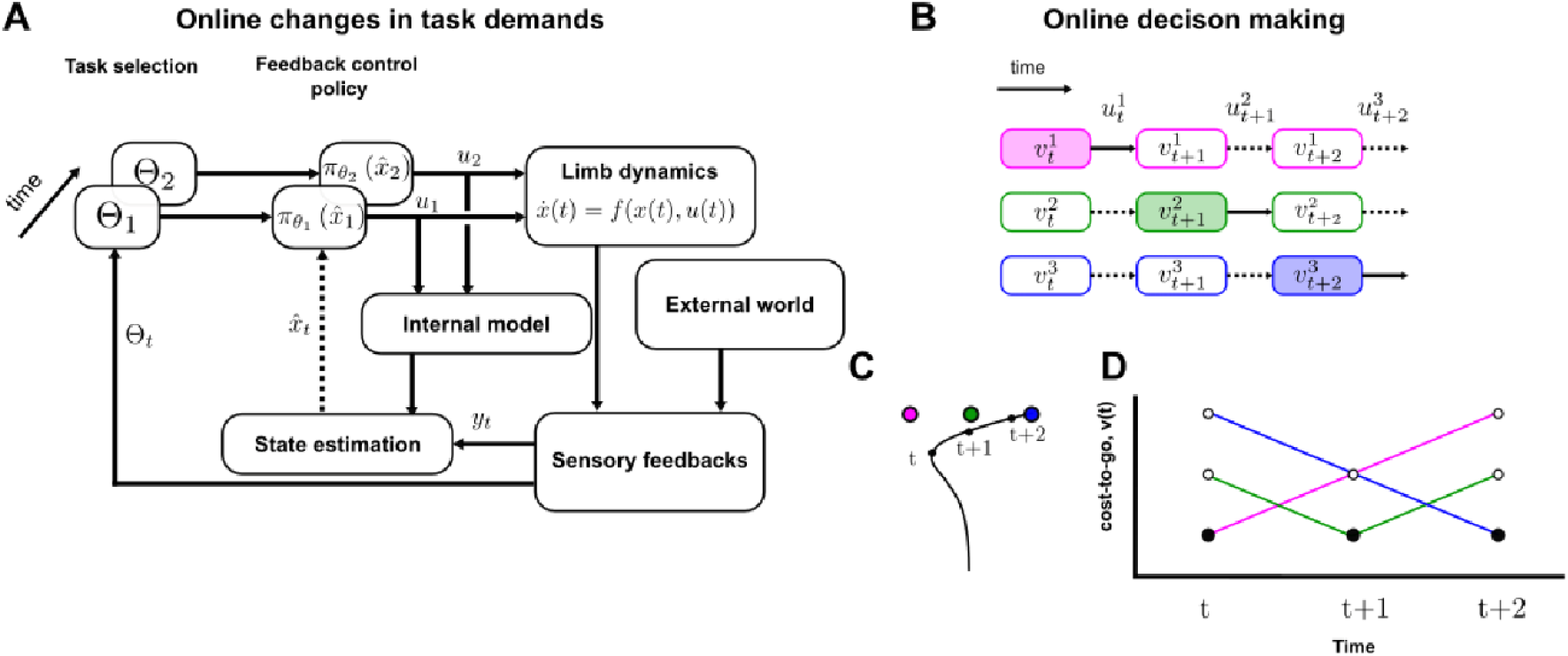
Model implementation. **A** Schematic representation of the computation of online changes in task demands. At each time step, the time-varying task parameters Θ_*t*_ are used to derive the optimal control policy 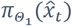 that is used to compute the motor command u_t_ (black arrow) which depends on the dynamical state estimate 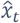 (dotted arrow) computed through dynamical Bayesian integration (state estimation). **B** Schematic representation of the implementation of online motor decisions in a three targets paradigm (see panel **C**). Each line schematized the time-varying cost-to-go functions associated with each option which are compared such that the one associated with the lowest value (see panel **D**) is selected (represented by the filled rectangle) and the corresponding motor command (full black arrow) is applied to the system for that very time step. **C** Representation of the different targets and an exemplar simulated hand trajectory (exaggerated case for illustration), the dots correspond to the time at which the decisions processes were considered. **D** Cost-to-go functions associated with the three targets evaluated at three different time points. The filled dots represent the minimum values that instructed the decision process, whenever these filled dots fall on a new color, it corresponds to an online change in target.

In order to validate this implementation, we simulated two different tasks that we previously conducted experimentally. The first task consisted of reaching movement towards a goal initially presented as a narrow or wide target, the main axis of which being orthogonal to the main reaching direction. During movement, the target width could instantaneously switch from narrow to wide or vice versa(17). The state vector penalty matrices (first term in equation (10)) for these two targets were defined such that the following equalities were verified, respectively for the narrow and wide targets

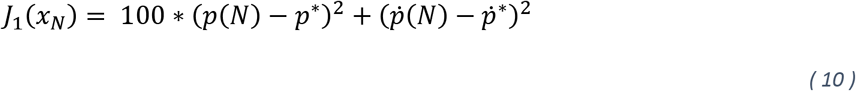

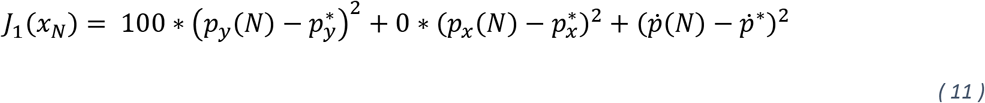

where the * superscript refers to the target state, *p*(*t*) is the position at time t, and the subscripts refer to the x- and y-coordinates. This means that there were no constraints on the end-point x-position of the wide target. The different gain values were arbitrarily selected such that the reaching behavior was qualitatively similar to what is observed experimentally. In addition to the changes in target structure, we induced mechanical step perturbations during movement by setting the value *F*_*ext*_ = 5*N* for the x-coordinate of this force.

The second task that we simulated to validate this implementation was the one presented in (18), which was similar to the previous one except that the target width could continuously change during movement. To model these phenomena, we defined the following time-varying state penalty matrices for the fast and slow conditions

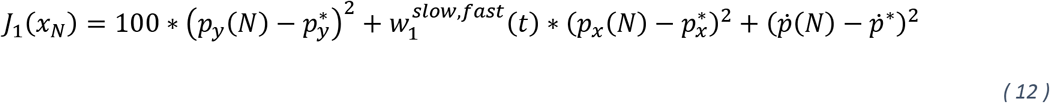

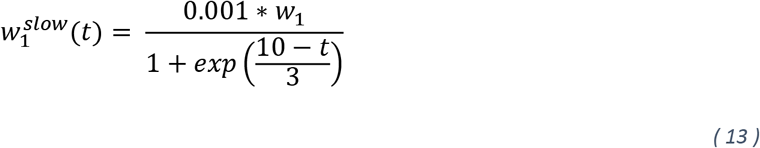

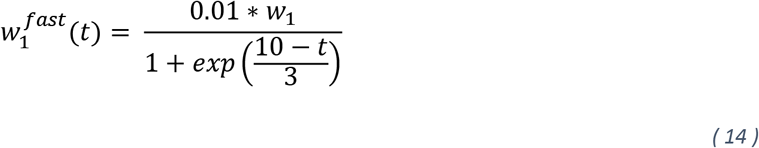

where *w*_1_=100. The time-dependency of these two parameters captures the continuous changes in target width.

In order to match the experimental finding that reported a delay of about 150ms between the change in target structure and the onset of differences observed in the muscle activity, we introduced a hard delay of 150ms (15 time-steps) to every modulation of target structure.

### Reward-dependent changes of mind

The classical formulation of the Optimal Feedback Control considers a single goal target which is used to derive the set of optimal feedback gains *L*_*t*_. In order to consider multiple potential targets during movement, Nashed and colleagues (25) proposed to leverage the cost-to-go function that appears in the dynamic programming resolution of the control problem (79). In this resolution, the optimal feedback gains are computed recursively thanks to the following backward recursion

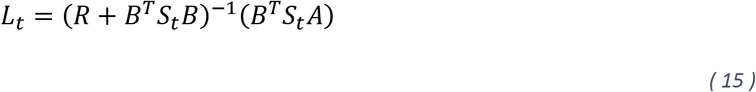

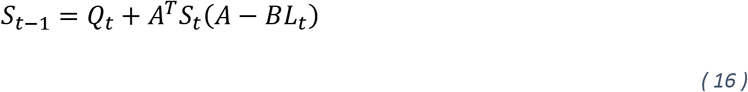

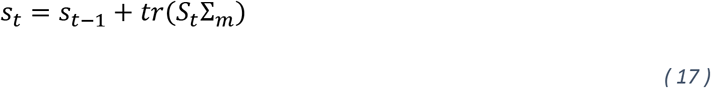

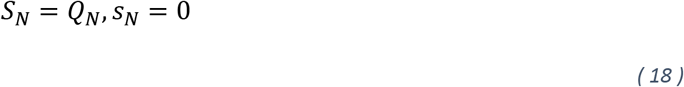

where *S*_*t*_ are real-valued matrices that capture the instantaneous penalty on the state vector, *s*_*t*_ is a scalar value involved in the computation of the cost-to-go function, and *tr*(.) denotes the trace operator. Under the optimal policy *π*, we define the estimated cost-to-go function as the remaining expected cost if the optimal policy is followed from the initial state 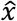 to the target state *x*^*^, this function writes as follows

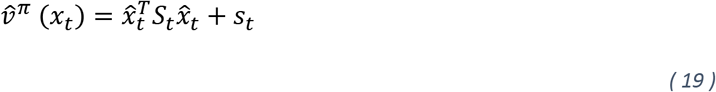

In theory, the cost-to-go is defined based on the true state vector *x*_*t*_ and on the impact of noise in the system, captured by the parameter *s*_*t*_. Here, however, the true state vector is not known from the point of view of the controller. Thus, because the best guess about the state is 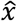, it is natural to consider that the best guess for the cost-to-go is the form in Equation (19). Note that the zero-mean Gaussian variable that is the error between the true and estimated state can be factored out of the first term of Equation (19), leading to an estimated cost-to-go equal to the true cost-to-go plus an error-term that influences *s*_*t*_. Similar developments apply if signal-dependent noise is considered, where error-related terms in the cost-to-go are already present.

To integrate multiple target states or alternative options for the movement, we computed the cost-to-go associated with each of these options similarly to what Nashed and colleagues proposed (25). In this study, all targets had the same reward; however, recent experimental reports demonstrated that reward could bias online motor decisions towards solutions that were not optimal if only sensorimotor factors were taken in isolation (20,21,23,47). Thus, it was necessary to alter the terminal value of the target to include these reward-related biases (*s*_*N*_).

Here we included explicit reward into the OFC formalism to allow reward-dependent biases in target selection as observed experimentally. In principle, Equation (19) represents an estimate of the total expected cost to accumulate from the current time step to the movement horizon. Clearly, an external reward can be taken in consideration by offsetting the value *s*_*t*_, which has a meaningful impact on the cost-to-go while leaving the recurrence and controller unchanged.

To simulate online motor decisions between alternative options, we computed the set of 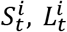 and 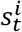 associated with each option *i*. Then, at each time step, the cost-to-go functions *v*^*i*^ associated with each option *i* are computed and compared, including rewards that were not related to the ongoing movement throughout the terms 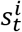, and the motor command associated with the smallest cost-to-go function is selected. Observe that if *v*^*i*^ < *v*^*j*^ for some time, and then suddenly *v*^*j*^ < *v*^*i*^ at some time, the corresponding control law (*u*^*j*^ instead of *u*^*i*^) effectively implements an online change in target. This implementation is novel in that it proposes a control framework to simulate reaching movements in presence of multiple targets potentially associated with different rewards. The basic premise is that, during movement, the controller continuously keeps track of the cost-to-go associated with each option by combining sensorimotor and cognitive factors and select the optimal action by comparing these different cost-to-go values. A schematic representation of this implementation is proposed in Figure 6B-D. Together, our implementation consists in a single feedback control policy that selects actions according to the current state of the body and environment to optimally solve the control problem.

We validated this implementation by comparing its predictions with experimental data collected in three experiments. The first experiment was similar to that reported in our previous study, the controller had to reach to any of three potential targets (21). For each target, we used the state penalty defined in Equation (10) and considered two different reward distributions: (i) all the targets had the same reward (*s*_0_ = 0) or (ii) the central target had a higher reward (*s*_0_ = 5. 10^−2^) than the other two (*s*_0_ = 0). The second and third experiments that we simulated to validate our model investigate reaching movements directed towards a redundant rectangular target that had a non-uniform reward distribution along its redundant axis, similarly to these used in (20,23). In the second experiment, we considered three different reward distributions along the redundant axis: (i) a symmetric distribution with higher rewards on both ends captured by equation (20) or an asymmetric distribution biased to the (ii) left or (iii) right, respectively captured by equations (21) and (22).

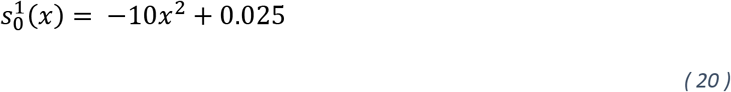

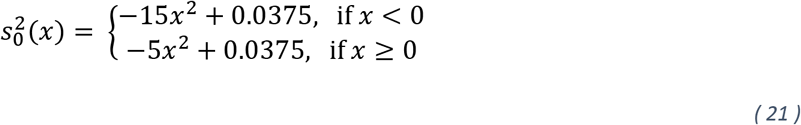

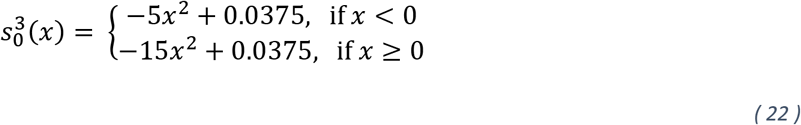

where *x* is the horizontal position (in cm) along the redundant axis, centered on zero. In all the simulations, we selected the reward values such that they could bias the cost-to-go function. Therefore, the selected values for *s*_0_ were of similar magnitude as the standard cost-to-go values.

### Data Analysis

Besides the simulation and characterization of human’s behavior in the various tasks mentioned above, we also used some of our previous experimental data to investigate whether the modulation of decision time predicted by our model was also present in the behavioral data. For this purpose, we investigated our previously collected dataset in which participants performed reaching movements towards multiple targets in presence of explicit target reward and perturbations of different magnitudes(21). More specifically, we investigated for a given reward distribution and perturbation amplitude the lateral hand deviation along the transverse axis to determine the onset of difference between the trials that reached the central and the lateral targets.

To extract the decision time, we used receiver operator curves (ROC) to determine the onset of change of mind in the kinematics. Briefly, the ROC curves compute the probability that two signals could be discriminated by an ideal observer. We compared trials with the same perturbation, the same reward distribution but with different final targets reached. The area under the ROC curve was calculated for the distribution of trials across participants and the onset of target-specific response was identified when the ROC curve exceeded 0.75 (80,81). The time reported in the Results section corresponds to that onset of target-specific differences.

## Acknowledgments

This work was supported by BELSPO (Belgian Federal Science Policy Office) and the European Space Agency. A.D.C. is supported by a K. Lisa Yang Integrative Computational Neuroscience (ICoN) Postdoctoral Fellowship. We thank Hari Kalidindi and Eric Wang for useful suggestions and comments.

## References

1. Lowrey CR, Nashed JY, Scott SH. Rapid and flexible whole body postural responses are evoked from perturbations to the upper limb during goal-directed reaching. J Neurophysiol. 1 mars 2017;117(3):1070–83.

2. Nashed JY, Crevecoeur F, Scott SH. Influence of the behavioral goal and environmental obstacles on rapid feedback responses. J Neurophysiol. 2012;108(4):999–1009.

3. Soechting JF. Effect of target size on spatial and temporal characteristics of a pointing movement in man. Exp Brain Res. 1984;54(1):121–32.

4. Codol O, Forgaard CJ, Galea JM, Gribble PL. Sensorimotor feedback loops are selectively sensitive to reward [Internet]. Neuroscience; 2021 sept [cité 8 mai 2022]. Disponible sur: http://biorxiv.org/lookup/doi/10.1101/2021.09.16.460659

5. Esteves PO, Oliveira LAS, Nogueira-Campos AA, Saunier G, Pozzo T, Oliveira JM, et al. Motor planning of goal-directed action is tuned by the emotional valence of the stimulus: A kinematic study. Sci Rep. 2016;6(March):1–7.

6. Summerside EM, Shadmehr R, Ahmed AA. Control of Movement Vigor of reaching movements : reward discounts the cost of effort. J Neurophysiol. 2018;119(6):2347–57.

7. Cross KP, Cluff T, Takei T, Scott SH. Visual Feedback Processing of the Limb Involves Two Distinct Phases. J Neurosci. 2019;39(34):6751–65.

8. Sabes PS, Jordan MI. Obstacle avoidance and a perturbation sensitivity model for motor planning. J Neurosci. 1997;17(18):7119–28.

9. Knill DC, Bondada A, Chhabra M. Flexible, Task-Dependent Use of Sensory Feedback to Control Hand Movements. J Neurosci. 2011;31(4):1219–37.

10. Georgopoulos AP, Kalaska JF, Massey JT. Spatial trajectories and reaction times of aimed movements: Effects of practice, uncertainty, and change in target location. J Neurophysiol. 1981;46(4):725–43.

11. Sarlegna FR, Mutha PK. The influence of visual target information on the online control of movements. Vision Res. 2015;110:144–54.

12. Crevecoeur F, Barrea A, Libouton X, Thonnard JL, Lefèvre P. Multisensory components of rapid motor responses to fingertip loading. J Neurophysiol. 1 juill 2017;118(1):331–43.

13. Forgaard C, Reschechtko S, Gribble PL, Pruszynski JA. Skin and muscle receptors shape coordinated fast feedback responses in the upper limb. Curr Opin Physiol. 2021;20:198–205.

14. Pruszynski JA, Johansson RS, Flanagan JR. A Rapid Tactile-Motor Reflex Automatically Guides Reaching toward Handheld Objects. Curr Biol. mars 2016;26(6):788–92.

15. Keyser J, Medendorp WP, Selen LPJ. Task-dependent vestibular feedback responses in reaching. J Neurophysiol. 2017;118(1):84–92.

16. Oostwoud Wijdenes L, van Beers RobJ, Medendorp WP. Vestibular modulation of visuomotor feedback gains in reaching. J Neurophysiol. 2019;122(3):947–57.

17. De Comite A, Crevecoeur F, Lefèvre P. Online modification of goal-directed control in human reaching movements. J Neurophysiol. 2021;125(5):1883–98.

18. De Comite A, Crevecoeur F, Lefèvre P. Continuous Tracking of Task Parameters Tunes Reaching Control Online. eNeuro. 2022;9(4):ENEURO.0055-22.2022

19. Cos I, Bélanger N, Cisek P. The influence of predicted arm biomechanics on decision making. J Neurophysiol. 2011;105(6):3022–33.

20. Cos I, Pezzulo G, Cisek P. Changes of mind after movement onset: a motor-state dependent decision-making process. eNeuro. 2021;8(6):ENEURO.0174.

21. De Comite A, Crevecoeur F, Lefèvre P. Reward-Dependent Selection of Feedback Gains Impacts Rapid Motor Decisions. eneuro. mars 2022;9(2):ENEURO.0439-21.2022.

22. Kurtzer I, Muraoka T, Singh T, Prasad M, Chauhan R, Adhami E. Reaching movements are automatically redirected to nearby options during target split. J Neurophysiol. 2020;124(4):10313–1028.

23. Martí-Marca A, Deco G, Cos I. Visual-reward driven changes of movement during action execution. Sci Rep. 2020;10(1):1–12.

24. Michalski J, Green AM, Cisek P. Reaching decisions during ongoing movements. J Neurophysiol. 2020;123(3):1090–102.

25. Nashed JY, Crevecoeur F, Scott SH. Rapid Online Selection between Multiple Motor Plans. J Neurosci. 2014;34(5):1769–80.

26. Scott SH. Optimal feedback control and the neural basis of volitional motor control. Nat Rev Neurosci. 2004;5(7):532–46.

27. Todorov E, Jordan MI. Optimal feedback control as a theory of motor coordination. Nat Neurosci. nov 2002;5(11):1226–35.

28. Todorov E. Optimality principles in sensorimotor control. Nat Neurosci. 2004;7(9):907–15.

29. Diedrichsen J. Optimal Task-Dependent Changes of Bimanual Feedback Control and Adaptation. Curr Biol. 2007;17(19):1675–9.

30. Diedrichsen J, Dowling N. Bimanual coordination as task-dependent linear control policies. Hum Mov Sci. 2009;28(3):334–47.

31. Izawa J, Shadmehr R. On-Line Processing of Uncertain Information in Visuomotor Control. J Neurosci. 2008;28(44):11360–8.

32. Omrani M, Diedrichsen J, Scott SH. Rapid feedback corrections during a bimanual postural task. J Neurophysiol. 2013;109(1):147–61.

33. Guigon E, Chafik O, Jarrassé N, Roby-Brami A. Experimental and theoretical study of velocity fluctuations during slow movements in humans. J Neurophysiol. 2019;121(2):715–27.

34. Lee JH. Model predictive control: Review of the three decades of development. Int J Control Autom Syst. 2011;9(3):415–24.

35. Ratcliff R, Smith PL, Brown SD, McKoon G. Diffusion Decision Model: Current Issues and History. Trends Cogn Sci. 1 avr 2016;20(4):260–81.

36. Liu D, Todorov E. Hierarchical optimal control of a 7-DOF arm model. In: 2009 IEEE Symposium on Adaptive Dynamic Programming and Reinforcement Learning [Internet]. Nashville, TN, USA: IEEE; 2009 [cité 11 janv 2022]. p. 50–7. Disponible sur: http://ieeexplore.ieee.org/document/4927525/

37. Todorov E, Li W, Pan X. From task parameters to motor synergies: A hierarchical framework for approximately optimal control of redundant manipulators. J Robot Syst. nov 2005;22(11):691–710.

38. Song S, Geyer H. A neural circuitry that emphasizes spinal feedback generates diverse behaviours of human locomotion. J Physiol. 2015;593(16):3493–511.

39. Mathew J, Crevecoeur F. Adaptive feedback control in human reaching adaptation to force fields. Front Hum Neurosci. 2021;15:742608.

40. Seethapathi N, Clark B, Srinivasan M. Exploration-based learning of a step to step controller predicts locomotor adaptation [Internet]. bioRxiv; 2021 [cité 11 mai 2022]. p. 2021.03.18.435986. Disponible sur: https://www.biorxiv.org/content/10.1101/2021.03.18.435986v1

41. García CE, Prett DM, Morari M. Model predictive control: Theory and practice—A survey. Automatica. mai 1989;25(3):335–48.

42. Dimitriou M, Wolpert DM, Franklin DW. The Temporal Evolution of Feedback Gains Rapidly Update to Task Demands. J Neurosci. 26 juin 2013;33(26):10898–909.

43. Cesonis J, Franklin DW. Time-to-target simplifies optimal control of visuomotor feedback responses. eNeuro. 2020;7(2):514–9.

44. Guigon E. A computational theory for the production of limb movements. Psychol Rev [Internet]. 12 août 2021 [cité 7 mars 2022]; Disponible sur: http://doi.apa.org/getdoi.cfm?doi=10.1037/rev0000323

45. Takagi A, Gomi H, Burdet E, Koike Y. A model predictive control strategy to regulate movements and interactions [Internet]. Neuroscience; 2022 août [cité 13 sept 2022]. Disponible sur: http://biorxiv.org/lookup/doi/10.1101/2022.08.24.505193

46. Codol O, Holland PJ, Manohar SG, Galea JM. Reward-based improvements in motor control are driven by multiple error-reducing mechanisms. J Neurosci. 2020;40(18):3604–20.

47. Cisek P. Making decisions through a distributed consensus. Curr Opin Neurobiol. 2012;22(6):927–36.

48. Trommershäuser J, Maloney LT, Landy MS. Statistical decision theory and trade-offs in the control of motor response. Spat Vis. 2003;16(3–4):255–75.

49. Bellman RE, Dreyfus SE. Applied dynamic programming. Princeton University Press; 1962.

50. Phillis YA. Controller Design of Systems with Multiplicative Noise. 1985;(10):1017–9.

51. Bertsekas D P. Reinforcement Learning and Optimal Control. Athena Scientifc; 2019.

52. Sutton RS, Barto AG. Reinforcement learning: An introduction, 2nd edition. Press MIT, éditeur. 2018.

53. Lillicrap TP, Hunt JJ, Pritze A, Heess N, Erez T, Y T, et al. Continuous control with deep reinforcement learning. arXiv. 2019;

54. Shadmehr R, Krakauer JW. A computational neuroanatomy for motor control. Exp Brain Res. 2008;185(3):359–81.

55. Mazzoni P, Hristova A, Krakauer JW. Why don’t we move faster ? Parkinson’s disease, movement vigor and implicit motivation. J Neurosci. 2007;27(27):7105–16.

56. Turner RS, Desmurget M. Basal ganglia contributions to motor control: A vigorous tutor. Curr Opin Neurobiol. 2010;70(6):704–16.

57. Shadmehr R, Reppert TR, Summerside EM, Yoon T, Ahmed AA. Movement Vigor as a Reflection of Subjective Economic Utility. Trends Neurosci. 2019;42(5):323–36.

58. Mink JW. THE BASAL GANGLIA: FOCUSED SELECTION AND INHIBITION OF COMPETING MOTOR PROGRAMS. Prog Neurobiol. 1 nov 1996;50(4):381–425.

59. Redgrave P, Prescott TJ, Gurney K. The basal ganglia: A vertebrate solution to the selection problem? Neuroscience. avr 1999;89(4):1009–23.

60. Dudman JT, Krakauer JW. The basal ganglia: from motor commands to the control of vigor. Curr Opin Neurobiol. 2016;37:158–66.

61. Thura D, Cisek P. The basal ganglia do not select reach targets but control the rugency of commitment. Neuron. 2017;95(5):991–3.

62. Bogacz R, Wagenmakers EJ, Forstmann BU, Niewenhuis S. The neural basis of speed-accuracy tradeoff. Trends Neurosci. 2010;33(1):10–6.

63. Thura D, Cisek P. Deliberation and commitment in the premotor cortex and primary motor cortex during dynamic decision making. Neuron. 2014;81(6):1401–16.

64. Cisek P, Kalaska JF. Neural Correlates of Reaching Decisions in Dorsal Premotor Cortex: Specification of Multiple Direction Choices and Final Selection of Action. Neuron. 3 mars 2005;45(5):801–14.

65. Dekleva BM, Ramkumar P, Wanda PA, Kording KP, Miller LE. Uncertainty leads to persistent effects on reach representations in dorsal premotor cortex. Frank MJ, éditeur. eLife. 15 juill 2016;5:e14316.

66. Manohar SG, Muhammed K, Fallon SJ, Husain M. Motivation dynamically increases noise resistane by internal feedback during movement. Neuropsychologia. 2019;123(4):19–29.

67. Berret B, Jean F. Why Don ‘ t We Move Slower? The Value of Time in the Neural Control of Action. 2016;36(4):1056–70.

68. Haith AM, Reppert TR, Shadmehr R. Evidence for Hyperbolic Temporal Discounting of Reward in Control of Movements. 2012;32(34):11727–36.

69. Shadmehr R, Orban de Xivry JJ, Xu-Wilson M, Shih T yu. Temporal Discounting of Reward and the Cost of Time in Motor Control. J Neurosci. 2010;30(31):10507–16.

70. Berret B, Castanier C, Bastide S, Deroche T. Vigour of self-paced reaching movement: cost of time and individual traits. Sci Rep. déc 2018;8(1):10655.

71. Berret B, Baud-Bovy G. Evidence for a cost of time in the invigoration of isometric reaching movements. J Neurophysiol. 2022;127(3):689–701.

72. Rigoux L, Guigon E. A Model of Reward- and Effort-Based Optimal Decision Making and Motor Control. Diedrichsen J, éditeur. PLoS Comput Biol. 4 oct 2012;8(10):e1002716.

73. Bian T, Wolpert DM, Jiang ZP. Model-free robust optimal feedback mechanisms of biological motor control. Neural Comput. 2020;32(3):562–95.

74. Crevecoeur F, Scott SH, Cluff T. Robust Control in Human Reaching Movements: A Model-Free Strategy to Compensate for Unpredictable Disturbances. J Neurosci. 2019;39(41):8135–48.

75. Brown IE, Loeb GE. Measured and modeled properties of mammalian skeletal muscle: IV. dynamics of activation and deactivation. J Muscle Res Cell Motil. 2000;21(1):33–47.

76. Crevecoeur F, McIntyre J, Thonnard JL, Lefèvre P. Movement stability under uncertain internal models of dynamics. J Neurophysiol. 2010;104(3):1301–13.

77. Crevecoeur F, Kurtzer I. Long-latency reflexes for inter-effector coordination reflect a continuous state feedback controller. J Neurophysiol. 2018;120(5):2466–83.

78. Česonis J, Franklin DW. Contextual cues are not unique for motor learning: Task-dependant switching of feedback controllers. PLOS Comput Biol. juin 2022;18(6):e1010192.

79. Kirk DE. Optimal control theory, an introduction. Dover Publications, Inc.; 2004.

80. Green DM, Swets JA. Signal detection theory and psychophysics. Oxford, England: John Wiley; 1966. xi, 455 p. (Signal detection theory and psychophysics).

81. Pruszynski JA, Kurtzer I, Scott SH. Rapid Motor Responses Are Appropriately Tuned to the Metrics of a Visuospatial Task. J Neurophysiol. 1 juill 2008;100(1):224–38.

